# No evidence of functional co-adaptation between clustered microRNAs

**DOI:** 10.1101/274811

**Authors:** Antonio Marco

**Author notes:** To whom correspondence should be addressed School of Biological Sciences, University of Essex, Wivenhoe Park, Colchester CO4 3SQ, United Kingdom.

## Abstract

A significant fraction of microRNA loci are organized in genomic clusters. The origin and evolutionary dynamics of these clusters have been extensively studied, although different authors have come to different conclusions. In a recent paper, it has been suggested that microRNAs in the same clusters evolve to target overlapping sets of genes. The authors interpret this as functional co-adaptation between clustered microRNAs. Here I reanalyze their results and I show that the observed overlap is mostly due to two factors: similarity between two seed sequences of a pair of clustered microRNAs, and the expected high number of common targets between pairs of microRNAs that have a large number of targets each. After correcting for these factors, I observed that clustered microRNAs from different microRNA families do not share more targets than expected by chance. During an exchange of correspondence and manuscripts, the authors of the original report acknowledged that the permutation methods they performed was not the method they described in their original paper. Here I show that the new permutation test proposed is biased and leads to systematic errors of the first kind, which will explain why the p-values reported were extremely (and unrealistically) low. I also discuss how to investigate the evolutionary dynamics of clustered microRNAs and their targets. In conclusion, there is no evidence of widespread functional co-adaptation between clustered microRNAs.

## INTRODUCTION

MicroRNAs are often clustered in the genome (Marco, Ninova, and Griffiths-Jones 2013). Whether these clusters have emerged by fusion of existing microRNAs, by tandem duplication, or by de novo emergence of new microRNAs near existing ones is still under debate, although a majority of clusters seem to have been originated after the emergence of a new microRNA next to an existing microRNA (Marco, Ninova, Ronshaugen, et al. 2013; Mohammed et al. 2013; Mohammed et al. 2014). In a recent paper, Wang et al. (2016) proposed that clustered microRNAs evolve to coordinately regulate functionally related genes. In their ‘functional co-adaptation’ model, clusters of microRNAs primarily emerge under the action of positive selection. Their model is an alternative to the ‘drift-draft’ model, which suggest that the maintenance of novel microRNAs in clusters is influenced by tight genetic linkage, and largely non-adaptive (Marco, Ninova, Ronshaugen, et al. 2013; Marco, Ninova, and Griffiths-Jones 2013). The ‘drift-draft’ model is based on the observation that a majority of microRNA clusters are formed during evolution by the random emergence of novel microRNAs near existing microRNA genes in the genome. Although functional co-adaptation may explain some observed patterns in specific microRNA clusters (see for instance (Chawla et al. 2016)), the evidence presented so far does not indicate that this is a widespread phenomenon. Here I show that the results presented by Wang et al (2016) do not support the conclusion that clustered microRNAs target overlapping sets of genes more than expected by chance, and therefore, there is no evidence of functional co-adaptation of clustered microRNAs.

In their paper, Wang et al (2016) first describe some evidence against the ‘drift-draft’ model. They expect that, if this model is correct, a higher fraction of microRNAs in introns should be conserved due to linkage effects. However, this is not a valid prediction: the ‘drift-draft’ model predicts that loss of microRNAs is reduced due to linkage, but also that the rate of *de novo* formation is increased (Marco, Ninova, Ronshaugen, et al. 2013). Therefore, one would expect both conserved and non-conserved (novel) microRNAs in introns. In other words, the different ratio of conserved/non-conserved microRNAs in introns with respect to intergeneric regions does not refute the ‘drift-draft’ model.

## RESULTS

### Clustered microRNAs do not target more common transcripts than expected by chance

Wang et al. (2016) reported that 1751 genes were targeted by at least two members of the same microRNA cluster. Reproducing their methodology (see Materials and Methods) I detected 1757 genes targeted by two or more microRNAs from the same cluster. Both estimates are very similar and the small differences may be due to slightly different parsing criteria. By performing a permutation analysis I found that this number is larger than expected by chance (10,000 permutations, p=0.0350, Figure 1A). However, a closer inspection to the results reveal that a large fraction of the common targets are attributed to only one cluster (*mir-183∼182*, 733 out of the 1757 targets, ∼42%), and that only 15 clusters had common targets between conserved microRNAs from different families, 5 of which were listed more than once, although the target count not added to the final number (Supplementary File 1). When we removed the *mir-183∼182* cluster from the analysis, the number of genes targeted by at least two clustered microRNAs (1024) is not significantly different to random expectations (p=0.2753, Figure 1A). Similarly, the number of genes targeted by three or more clustered microRNAs (77) was not significantly high (p=0.9887). In conclusion, the apparent overlap among the targets of clustered microRNAs is due to only one cluster: *mir- 183∼182*.

**Figure 1.**
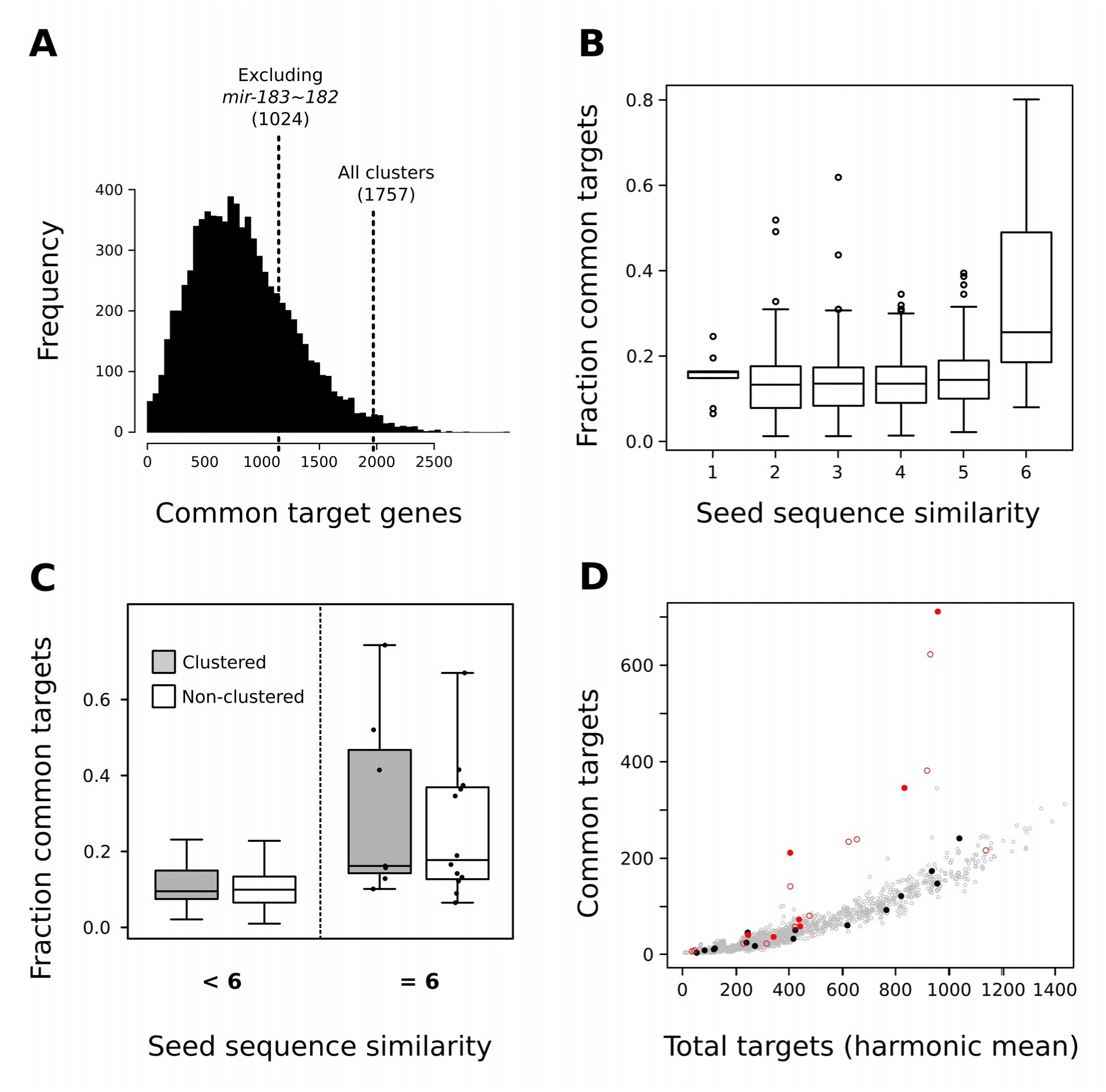
Common targets between clustered and non-clustered microRNAs. (A) Permutation analysis of common targets between clustered microRNAs. Each of the 10,000 permutations reported the number of common targets between at least two microRNAs from the same cluster. The observed values for all clusters, and for all but one outlier (*mir-183∼182*) are indicated with vertical dashed lines. (B) Fraction of common targets (common targets divided the minimum number of individual targets) for all microRNAs in this study, binned by the seed sequence similarity. (C) Fraction of common targets for clustered and non-clustered pairs of microRNAs, for those with a seed similarity less than 6 (left panel) or 6 (right panel). (D) Number of common targets between pairs of microRNAs versus the harmonic mean of the individual number of targets for the pair. Filled circles represent clustered microRNAs, gray circles are unclustered microRNAs, and red circles are pairs of microRNAs with a seed similarity of 6.

Why does this cluster have a large number of overlapping targets? The *mir-183∼182* cluster host three microRNA families with targets, yet the majority of the common targets reported (711) are found between two microRNA families: miR-183-5p and mir-96-5p/1271-5p. Both families have very similar seed sequences (AUGGCAC and UUGGCAC), which differ in only the first nucleotide. Similar seeds are more likely to have overlapping sets of targets randomly. In fact, when two microRNA families share 6 out of the 7 nucleotides that define the seed sequence, the number of common targets is significantly high (Figure 1B). When binning the data into pairs of microRNAs that are found clustered and those unclustered, accounting for those pairs of families with 6 common nucleotides, we observed that the differences between clustered and non-clustered microRNAs are not significant (p=0.235; Figure 1C)^1^. In conclusion, the observed overlap between targets in some clustered microRNAs is actually the random consequence of the similarity between their seed sequences, and is not associated to whether the microRNAs are clustered or not. Interestingly, mir-183 and mir-96 have similar seed sequences but they are probably not paralogs. One possibility that directional selection have change the seed sequences of one (or both) microRNAs during evolution. However, both seed sequences have been conserved since their origin (Supplementary Figure 1A) and, therefore, there is no evidence of substitutions happening in the seed of these microRNAs for the last 600 million years.

From these results it is also evident that clusters sharing a large number of targets are those whose individual microRNAs target a large number of transcripts. This is expected from a probabilistic point of view since, the more targets two microRNAs have, the more the chances that some of them will be shared. In Figure 1D I plot the number of overlapping targets as a function of the harmonic mean of the number of targets of each microRNA in a pair. A few outliers to a near perfect correlation correspond to microRNA family pairs with similar seed sequences (red circles). There is not a clear difference between clustered and non-clustered microRNAs (Figure 1D).

### The permutation methods: the main source of discrepancy

After a first version of this manuscript was published, Wang et al. (2018, doi:10.1101/313817) responded by admitting that **the permutation method they used was different to that described in their original report** which, in my opinion, invalidates the findings reported in their previous manuscript. In Wang et al (2016) they wrote: “in the permutation analysis, we only randomly shuffled the locations of miRNAs”. However, they claim in Wang et al. (2018) that: “we first shuffled the co-expressed seed:target pairing […] and then we tested how many genes were targeted by at least two miRNAs (with distinct seed) in the same cluster”. The two methods will lead to very different results, as I explain in detail below.

The literature on permutation methods (so-called exact tests) is vast (Efron 1982; Good 2006). During a permutation test we rearrange the covariates, usually by shuffling the labels or the frequencies in the test (Sokal and Rohlf 1995). This is done over the condition to be tested, in our case, over the ‘clustering’ condition. That is, to test whether clustered microRNAs have more (or less) common targets than expected by chance, we permute the microRNA ‘labels’ across the existing loci (clustered and unclustered), keeping all other conditions unaltered. This is indeed the method used here, and also originally described by Wang et al (2016). Interestingly, the same idea of shuffling the miRNA labels and not the interactions was described in another work from the same lab (Luo et al. 2018).

Why we should not shuffle the microRNA-transcript interactions? Because by permuting over interactions, we are not testing the effect of clusters but whether some genes have more (or less) interactions than others, which is obviously true (due to variation in length and composition of 3’UTRs) and does not add anything to the clustering problem. I will leave the verbal argument and show a clear example. Figure 2A shows a cartoon of a hypothetical regulatory network. Two microRNAs are clustered and another one is unclustered. All microRNAs target the same transcript ‘a’ (that is, ‘a’ is a promiscuous target). Additionally, the unclustered microRNAs target a second transcript ‘b’. Therefore, ‘a’ is not more likely to be targeted by a clustered than by an unclustered microRNA. If we permute the microRNA labels keeping everything else constant, all permutations will report that there is one gene targeted by two clustered microRNAs and, therefore, the probability of observing that there is one gene targeted by two clustered microRNAs in Figure 2A is obviously 1. In other words, the observed common target will not be statistically different to our random expectation, and the test will report a p-value of 1. Alternatively, if we shuffle the interactions in Figure 2A, the probability of observing by chance that one gene targeted by two clustered microRNAs is 0.5 (which can be found by the total enumeration of cases). That is, a p-value computed from this distribution will be half the expected value under the null hypothesis.

**Figure 2.**
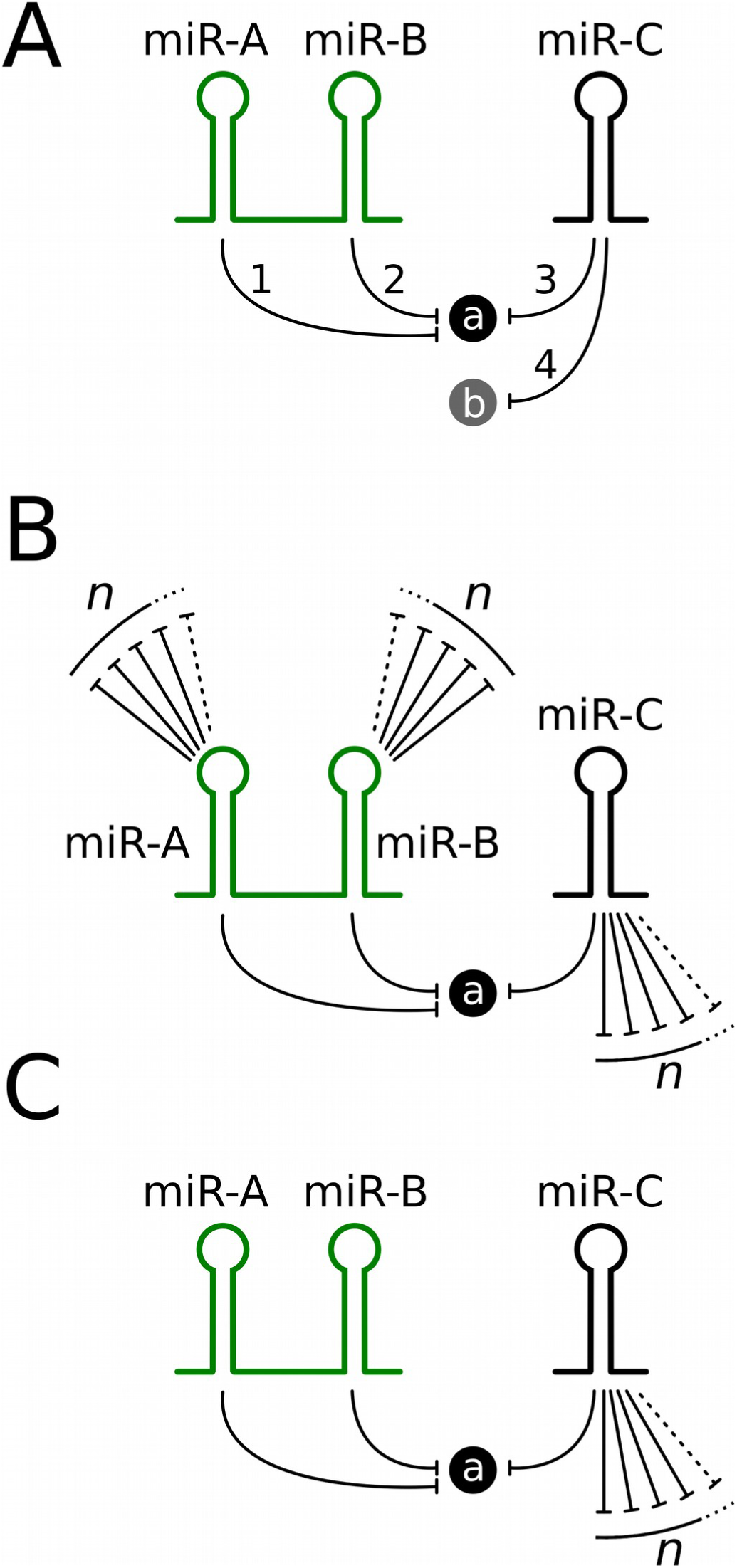
Examples of simple microRNA:transcript regulatory networks. Three simple networks used to illustrate the effect of the permutation method on the computation of p-values.

If we allow all microRNAs to have a number *n* of additional interactions (Figure 2B), the underestimation of p-values is also evident. The probability in Figure 2B that a permutation reports a common target between two clustered microRNAs converges to 0.44 (Wang 2018, doi:10.1101/313817, version April 14, 2019)^2^, whilst under appropriate statistical testing this probability should be 1. Cases in which the number of target differ between element of the regulatory network produce even smaller p-values by chance. For instance, in Figure 2B, using standard combinatorics (Biggs 2002), we can compute *p* as: 

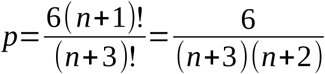

Thus, for n=9, this permutation method will detect a significant enrichment of clustered microRNAs with common targets with a p-value of 0.045.

Now, let’s explore the technical aspects of the significance test. According to standard statistical inference theory, when the null hypothesis is true, the p-values from a statistical test should follow a uniform distribution between [0,1]. To investigate the distribution of p-values for the two shuffling methods proposed, I randomly constructed 10,000 microRNA sets, with the same clustering configuration as that analyzed in the papers, but restricting the shuffling such that clusters of microRNAs will be composed only by microRNAs that are not clustered in the human genome. That is, for all synthetic sets, the null hypothesis ‘clustered microRNAs share as many common targets than non-clustered microRNAs’ is true. For each set I computed the p-value by shuffling the microRNA loci (as I propose) for both the one- and the two-tailed versions of the test. This method produces p-values that are distributed uniformly (Figure 3A,B). Then, I computed the p-values resulting from permuting the microRNA:gene interactions as Wang et al. (2018) suggest. The p-values in this case are L-shaped and U-shaped distributed for the two- and the one-tailed versions of the test respectively. Indeed, by evaluating the sets for which the null hypothesis is true, the test proposed by Wang et al. (2018) significantly rejected the null hypothesis in the vast majority of cases. This most likely explains why all the p-values reported by Wang et al. (2016,2018) were extremely low: almost any cluster configuration explored will show significant target overlaps.

**Figure 3.**
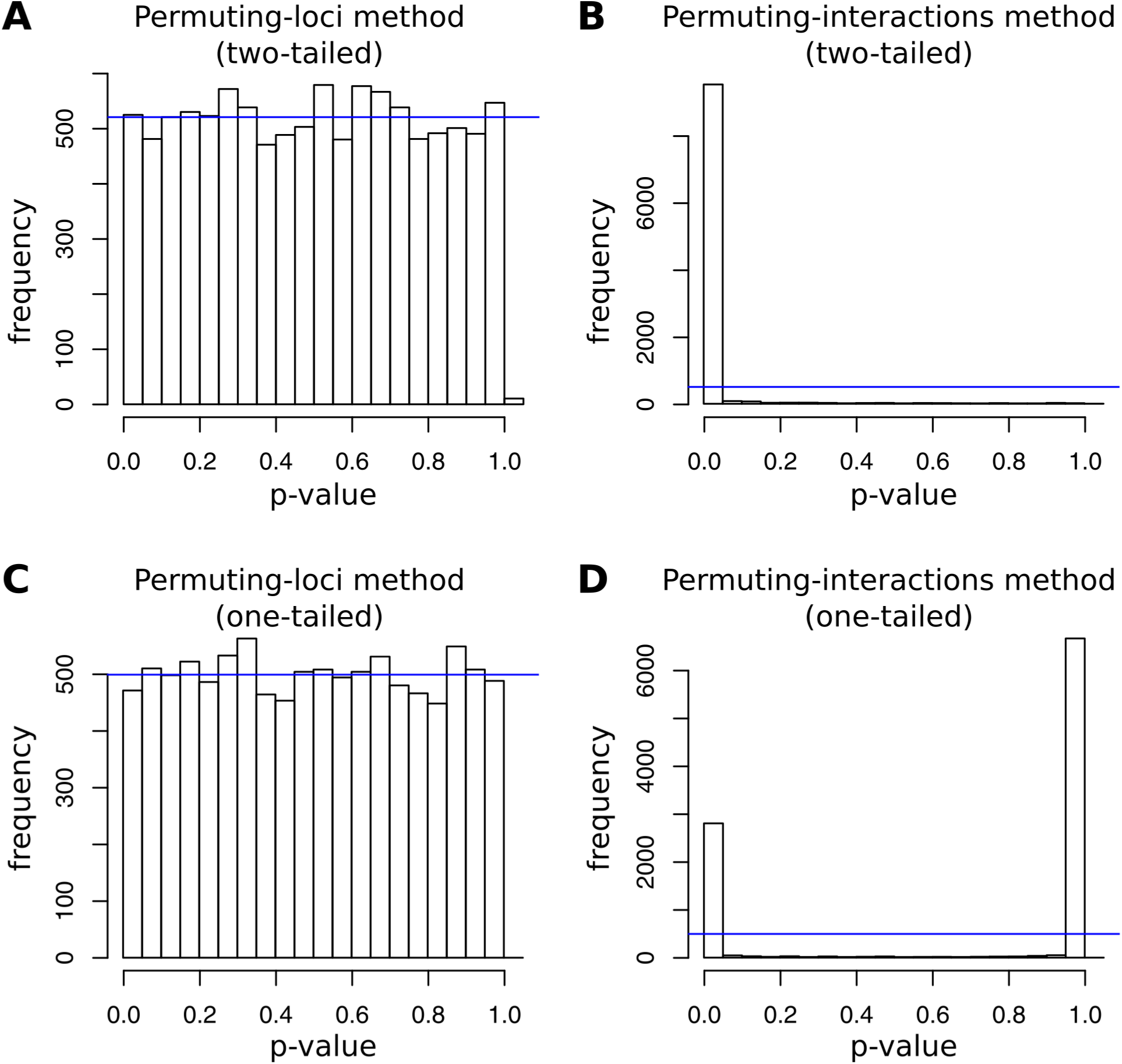
Distribution of p-values from different permutation tests. A) Permuting loci, two-tailed test; B) Permuting interactions, two-tailed test; C) Permuting loci, one-tailed test; D) Permuting interactions, one-tailed test.

In summary, all statistical tests performed by Wang et al. (2016,2018) based on the permutation of interactions yielded unrealistically low p-values. Thus, the main results of these manuscripts are not reproducible.

## DISCUSSION

Early reviews speculated that clustered microRNAs may target common genes (Bartel 2004; Kim and Nam 2006). Indeed, the pioneering analysis by (Grün et al. 2005) showed evidence that, in *Drosophila*, some clustered microRNAs tend to have common targets. However, when using the latest available data, this observation is not statistically significant (p=0.2569; see Methods). Some specific cases have been suggested: non-paralogous microRNAs in the mir-106b∼25 cluster have a few CDK inhibitors as common targets (Kim et al. 2009); four microRNAs from the mir-379∼410 cluster may target the gene PUM2 during dendrite differentiation (Fiore et al. 2009); and members of the miR-23a∼24-2 cluster have common targets (Chhabra et al. 2010). In all these examples, microRNAs are highly conserved and show no nucleotide differences in their seed regions in all studied species (Supplementary Figure 1B-D). This indicates that observed changes in the microRNA-mRNA regulatory networks occur mostly at the level of target sites rather than in microRNAs loci (see further discussion below). Whether functional co-adaptation operated among the members during the early stages of cluster formation cannot be determined using the current methodology. But the fact that all examples studied are highly conserved, leaves little room at this moment to this possibility.

Wang et al. (2016) also claimed that microRNAs in clusters may be involved in common biological pathways. In the pathway enrichment analysis (in their supplementary Table S9), a large majority of processes enriched for target sites of clustered microRNAs correspond to sites from the mir-183∼182 cluster or other clusters with similar seed sequences among their members. After removing these clusters the pathway enrichment is explained by two seed families from the cluster mir-17∼92a: miR-17-5p/20-5p/93-5p/106-5p/519-3p and miR-19-3p. As with the other examples discussed so far, the seed sequences of miR-17-5p and miR-19a-3p are highly conserved (accession numbers MIPF0000001 and MIPF0000011 in http://mirbase.org). This further support that changes in target sites, rather than in microRNA loci, may explain the observed functional overlap between the targets of clustered microRNAs.

Interestingly, in a recent update of Wang et al’s response, the authors suggested a new analysis in which target sites are selected arbitrarily (cumulative weighted context++ score < −0.3), which means that target conservation is not taken into account. In this analysis it seems that clustered microRNAs tend to target more common targets than expected by chance (marginal significance yet detectable). This, together with the fact that the overlap is not significant when accounting for conserved targets, and that analyzed clustered microRNAs are actually conserved for millions of years, further supports that the changes determining the microRNA:target evolutionary dynamics are mostly happening at the target sites rather than at the microRNA seed sequences. In other words, this novel analysis does not prove adaptive selection of clustered microRNAs but evolutionary changes at the target sites, whose evolutionary mechanism is yet to be elucidated.

How microRNA clusters emerge is a relatively known mechanism where discrepancies between authors are mostly around specific details (Marco, Ninova, Ronshaugen, et al. 2013; Mohammed et al. 2013; Mohammed et al. 2014). However, the impact of clustered microRNAs in the evolution of target sites is a largely unexplored research area. Perhaps is time now to focus more on the targets, rather than in the the microRNAs, to understand why functionally related genes are sometimes targeted by clustered microRNAs. In summary, the model proposed by Wang et al. (2016) is an interesting hypothesis that provides a mechanistic explanation to microRNA clusters. Indeed, their functional co-adaptation model may be tested as new datasets and/or methodological frameworks become available. However, under the current available data, there is no statistical evidence that clustered microRNAs have evolved to target common genes, and microRNAs targeting functionally related genes do not show evidence of adaptation. In conclusion, there is no evidence of widespread functional co-adaptation between clustered microRNAs.

## METHODS

The datasets and protocols to predict microRNA targets were as described in Wang et al. (2016). Target predictions and microRNA families were downloaded from TargetScanHuman, release 7.1 (Agarwal et al. 2015). The cutoff to select target in human genes was P_ct_ ≥ 0.5 and the microRNA families were classified in two groups: conserved among animals and broadly conserved among vertebrates. MicroRNA expression data was extracted from miRBase release 21 (Kozomara and Griffiths-Jones 2014) for the experimental data from Meunier et al. (Meunier et al. 2013). Only microRNAs with 50 or more reads were further considered. Transcript expression data was obtained from Brawand et al (2011). Only co-expressed microRNA-target pairs were analyzed. Two microRNAs were considered to be in the same clusters if they were less than 10000 nucleotides apart from each other in the human genome GRCh38, as indexed in miRBase. Pairs of microRNA families coming from the same precursor were not taken into account (which may, in some cases, target overlapping sets of transcripts [Marco et al. 2012]). Overlapping microRNA clusters were not considered in the final counts, but they are listed in the supplementary files. For the permutation analysis, only the location of the microRNAs was permuted, keeping the structure of the clusters; the common targets were counted in 10,000 independent random replicates. Seed sequence similarity is defined as the longest common substring. MicroRNA targets from PicTar were retrieved from UCSC Table Browser (Genome: dm2; table: picTarMiRNAS1), *Drosophila* microRNA clusters are those described by Grün et al. (2005), and the statistical significance was assessed by randomly permuting the microRNA loci across clusters (10,000 replicates). All analysis performed can be reproduced by running a collection of scripts provided in a UNIX operating system (https://doi.org/10.6084/m9.figshare.6165722).

## ACKNOWLEDGEMENTS

I thank the authors of Wang et al 2016 for comments on the manuscript and spotting a bug in an earlier version of the code. This work was supported by the Wellcome Trust [grant number 200585/ Z/16/Z].

## Footnotes

1 In their response, Wang et al claimed that when computing the harmonic mean instead of the fraction of common targets, the results are different to those here reported. However, when evaluating their source code the program counts several times the same interaction. When this is corrected the results are very similar to those here reported.

2 Wang et al identified an error I made in the previous version in which I provided a formula for a p-value for a different network configuration. Yet, their corrected p-value estimation is still below 1.

## Notes

#### Summary of Updates

Small errors corrected and two notes added to clarify the discrepancies between this manuscript and a response. NOTE FROM PREVIOUS REVISIONS: - In this revision I corrected a bug in one of the programs and updated the manuscript accordingly. The results remained unaltered. Additionally I added some discussion on a topic raised by other authors regarding non-conserved targets, which turned out to support the model I proposed earlier. - The previous version has a response in which the authors acknowledge that a different method was used in their original analysis. In this version I reproduce the new methodology and show why it leads to incorrect p-values.

