## Supplementary Figure 1 for "No evidence of functional co-adaptation between clustered microRNAs"

**Supplementary Figure 1. Seed sequence conservation in clustered microRNA familes.** Sequences were downloaded from miRBase (http://mirbase.org). Red areas indicate the seed region of the mature sequence. Alignments represent mature sequences with a reported common target from the cluster: A) mir-183~182; B) mir-106b~25; C) miR-23a~24-2; D) mir-379~410. Species code as defined in miRBase.

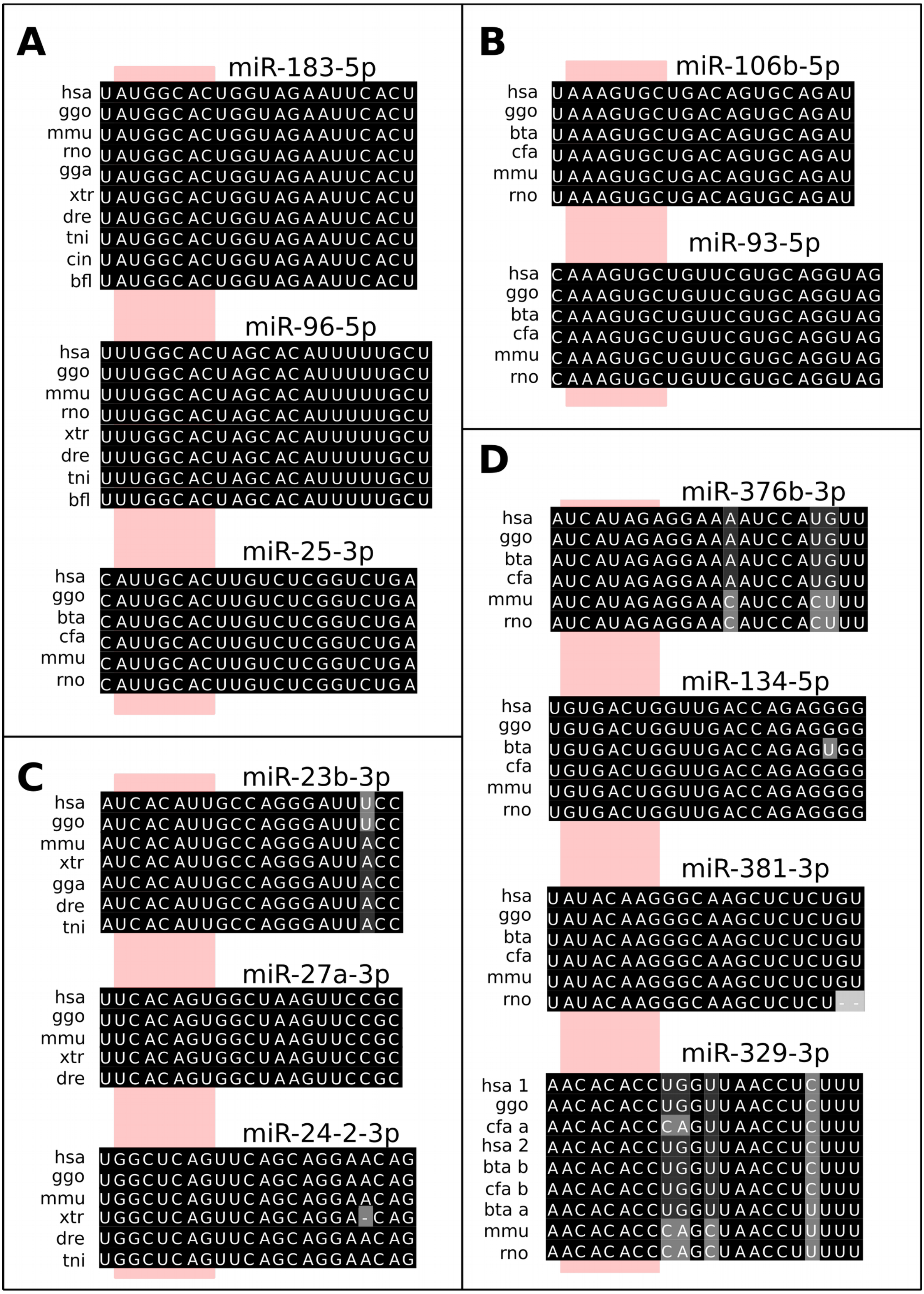
